# A flexible electronic strain sensor for the real-time monitoring of tumor progression

**DOI:** 10.1101/2021.09.16.460551

**Authors:** Alex Abramson, Carmel T. Chan, Yasser Khan, Alana Mermin-Bunnell, Naoji Matsuhisa, Robyn Fong, Rohan Shad, William Hiesinger, Parag Mallick, Sanjiv Sam Gambhir, Zhenan Bao

**Affiliations:** Department of Chemical Engineering, Stanford University, Stanford, CA, 94305; Department of Radiology, Stanford University, Stanford, CA, 94305; Molecular Imaging Program at Stanford (MIPS) and Bio-X Program, Stanford University, Stanford, CA, 94305; Department of Bioengineering, Stanford University, Stanford, CA, 94305; Department of Cardiothoracic Surgery, Stanford University School of Medicine, Stanford, CA; Department of Medicine, Stanford University, Stanford, CA, 94305; Canary Center at Stanford for Cancer Early Detection, Stanford University, Stanford, CA, 94305

**Author notes:** Deceased (Sanjiv Sam Gambhir). Department of Electronics and Electrical Engineering, Keio University, Yokohama, Kanagawa 223-8522, Japan (Naoji Matsuhisa).

## Abstract

Healthcare professionals and scientists utilize tumor shrinkage as a key metric to establish the efficacy of cancer treatments. However, current measurement tools such as CT scanners and calipers only provide brief snapshots of the dynamic geometric changes occurring in vivo, and they are unable to characterize the continuous micrometer-scale volumetric transformations transpiring at minute timescales. Here we present a stretchable electronic strain sensor, with a 10-micron scale resolution, capable of continuously monitoring tumor volume progression in real-time. In mouse models with subcutaneously implanted lung cancer or B-cell lymphoma tumors our sensors discerned a significant change in the tumor volumes of treated mice within 5 hours after small molecule therapy or immunotherapy initiation. Histology, caliper measurements, and luminescence imaging over a one-week treatment period validated the data from the continuous sensor. We anticipate that real-time tumor progression datasets could help expedite and automate the process of screening cancer therapies in vivo.

## Introduction

When treating cancer, time is of immense value (*1*). However, no tools currently exist to read out in vivo tumor progression in real-time, even in pre-clinical models. A lack of understanding around in vivo temporal tumor progression inhibits rapid clinical decision-making (*2*). Currently, many clinicians discern cancer treatment efficacies based on gross tumor regression measured by sequential CT scans (*3*–*5*). Yet, due to the months-long delays between treatments and the extended time periods required to discern a biologically significant change between scans, patient mortality rates can reach 20% before healthcare professionals fully evaluate the efficacy of a treatment in a given patient (*1, 6, 7*). Preclinical animal testing for cancer therapeutics suffers similarly slow readout times, often requiring days, weeks, or even months before a treatment can be deemed effective (*8*). While tumor cell culture and organoid experiments can demonstrate treatment responses from patient cells on the order of hours (*9, 10*), the same timetables are unable to be replicated in animal models (*11*) due to a lack of measurement tools capable of precise, continuous monitoring. Mechanical instruments such as calipers possess measurement errors of up to 20% (*12–14*), preventing the detection of micrometer-scale tumor volume dynamics. Moreover, imaging tools such as CT scanners and Bioluminescence Imagers possess cost, usage, and safety barriers that eliminate the possibility of continuous measurement (*15*). A device capable of continuously monitoring tumor progression in vivo could provide insights into the dynamics of tumor growth and shrinkage that allow for faster and more efficacious therapeutic regimens.

Here, we present an elastomeric-electronic tumor volume sensor capable of reading out cancer treatment efficacy studies within hours after therapy initiation. Utilizing advances in flexible electronic materials (*16*–*21*), we designed a conformal strain sensor that continuously measures, records, and broadcasts tumor volume changes at a 10 μm scale resolution, approximately the size of a single cell. This sensor achieves three main advances over other common tumor measurement tools such as calipers and imagers. First, because the sensor remains in place over the entire measurement period and takes measurements every five minutes, it is possible to generate a four-dimensional, time dependent dataset that eliminates the need for any guesswork on measurement timing. Second, the sensor possesses the capability of detecting size changes that fall within the error of caliper and imaging measurements, allowing for more precise readouts that catch smaller tumor volume changes. Third, the sensor is entirely autonomous. Therefore, using it reduces the costs and labor associated with performing measurements, and it enables direct data comparisons between operators. It therefore enables fast, cheap, large-scale preclinical drug discovery testing setups. We call our technology FAST, which stands for Flexible Autonomous Sensors measuring Tumor volume progression.

## Results

### Designing a strain sensor for measuring tumor volume progression

Our wireless FAST technology for real-time monitoring of tumor size progression can be applied to tumors on or near the skin (Fig 1a). The sensor is fabricated by depositing a 50 nm layer of gold on top of a drop casted layer of styrene-ethylene-butylene-styrene (SEBS), and it can be easily scaled up for mass manufacturing (see Supplementary Text and Supplementary Figure S1). Because the sensor is fully flexible and stretchable, it readily expands or shrinks with the tumor as it progresses. Compared to other homogenous sensors that increase linearly with strain, the resistance in this sensor rises exponentially as strain grows, as explained through percolation theory; when strain is applied, microcracks in the gold layer lose contact with each other, increasing the tortuosity of the electron path length through the sensor (Fig 1b). The relative change in resistance in the sensor spans two orders of magnitude as it is stretched from 0% to 75% strain and can detect changes down to a 10 μm scale resolution (Fig 1c,d). At 100% strain, the electrical connection between the two ends of the sensor breaks; however, the sensor can stretch to over 200% strain before the SEBS ruptures (see Supplementary Figure S2), and it is able to regain an electrical connection when the sensor returns to a lower strain. By changing the thickness of the SEBS layer (Fig. 1e and see Supplementary Figure S2), it is possible to increase the stress that can be applied to the sensor before it ruptures.

**Figure 1:**
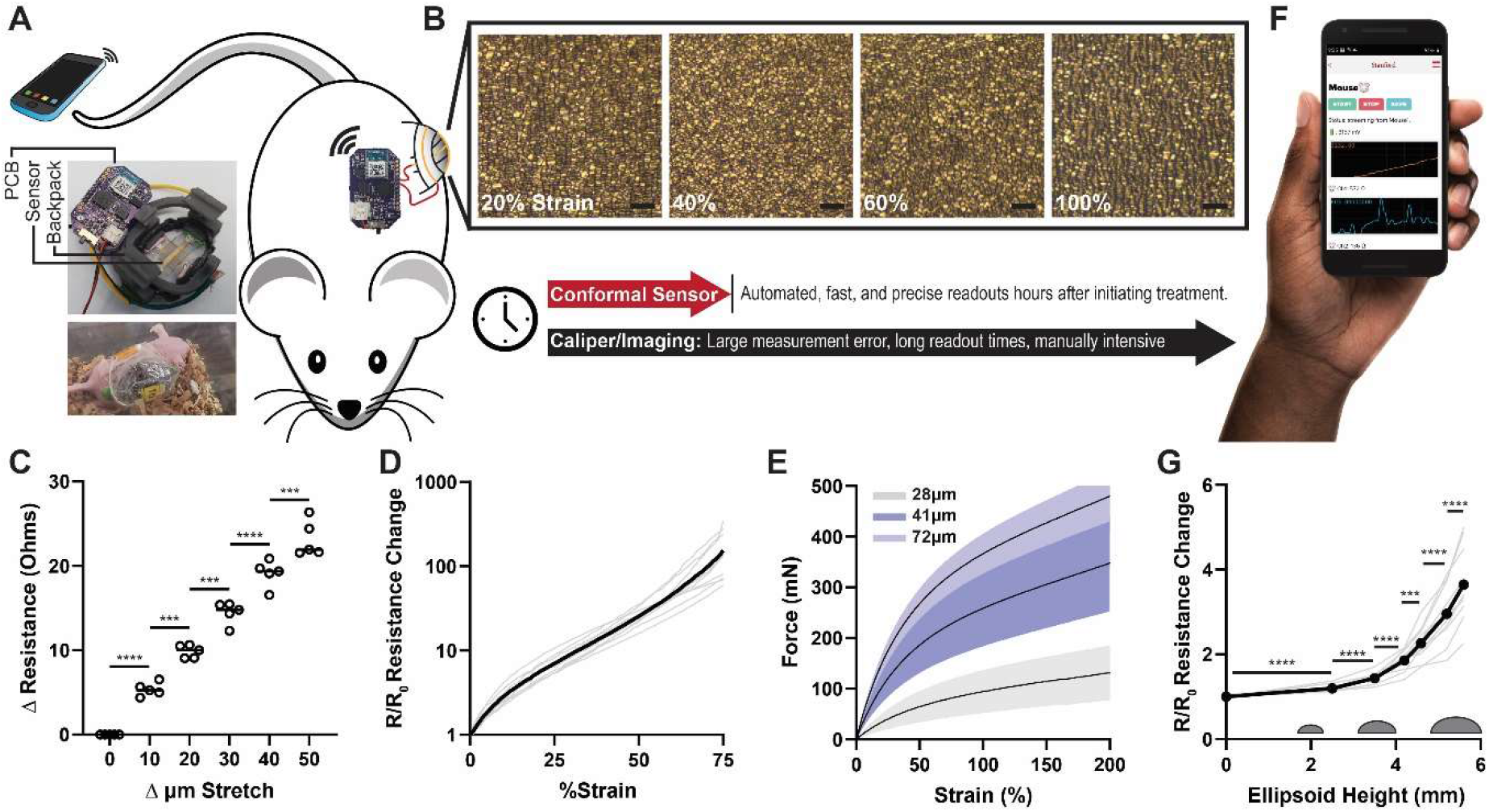
Flexible Autonomous Sensors measuring Tumor volume progression (FAST). (A) Schematic images of the FAST technology containing a printed circuit board (PCB), stretchable strain sensors, and a backpack to hold the sensor on the mouse. (B) Light microscopy images of a cracked gold strain sensor at varying strains. (SB=20 μm). (C) Recorded resistance changes in cracked gold strain sensors as they are stretched in 10 μm increments from an initial pre-strain of 50% (n=5, Induvial Data Points, Line = Median; One way ANOVA with Tukey’s multiple comparisons test). (D) Recorded fold change in resistance of cracked gold strain sensors as they are strained from 0% to 75% strain (n=10, Individual Curves, Bold Line = Average). (E) Force required to strain cracked gold sensors of varying SEBS substrate thicknesses (n=12 or 13, Line= Average ± SD). (F) Rendered screen shot from a custom cell phone app recording the resistance change in the strain sensor on a mouse possessing a treated tumor. (G) Recorded fold change in resistance of FAST as it measures increasing sizes of 3D printed replica ellipsoid-shaped tumors with volumes comparable to in vivo mouse tumors (n=10, Individual Curves, Bold Line = Average; ratio paired t test). (*p<0.05; **p<0.01; ***p<0.001; ****p<0.0001).

A custom designed printed circuit board and cell phone app enable live and historical sensor readouts with the press of a button (Fig 1f and see supplementary figure S3). To read out the sensor, it is placed in series with a known resistor on the board, and a known voltage is applied across the circuit. The voltage drop over the known resistor is amplified by an instrumentation amplifier, converted to a digital signal, and read out by an analog-to-digital converter (ADC) of a microcontroller. To read out resistances between 300-60,000Ω accurately and precisely, the circuit board applies three different voltage biases through the resistive sensor and chooses the most accurate reading depending on the sensor’s resistance. Moreover, the circuit board takes 32 consecutive measurements and reports the median readout, ensuring that the data is not contaminated by slight movements. We measured the error in sensor readout to be 1-2%, as calculated through measurements of known resistors (see Supplementary Fig S3). The assembled device can read out measurements continuously every 5 minutes for >24 hours on a 150 mAh battery. Further optimization of the machine code would increase the battery life closer to the theoretical maximum of measurements once per hour for >10 days.

We designed a 3D printed housing mechanism for FAST to ensure that the sensor and PCB fit comfortably on the mouse and accurately record tumor volume progression (Fig 1a and Supplementary Fig S1). The housing possesses a flexible base capable of conforming to the mouse’s skin as well as rigid rods that ensure the ends of the sensors remain fixed in place. Fixing the ends of the sensors to rigid components, rather than placing them directly on flexible skin, allows us to calculate the sensor’s change in strain attributed to tumor growth without the additional convoluting factor of skin displacement. The sensors themselves are pre-stretched up to 50%, enabling us to accurately read out both growth and shrinkage events within the device’s most sensitive strain range. To characterize the assembled device’s ability to discern volume variations in shapes in vitro, we measured the sensor’s output when placed on top of 3D printed model tumors (Fig 1g). The sensors recorded significant changes in readouts for objects as small as 2.5 mm in height and as large as 5.6 mm in height. Changing the initial strain on the sensor allows for the measurement of larger objects as well. We provide a method for calculating the size of an object using the sensors readouts from our assembled device in the supplementary material.

### Continuously Tracking Tumor Progression In Vivo

In vivo testing in two cancer models demonstrated that FAST detected tumor growth and shrinkage in mice faster than and with a comparable or greater accuracy to calipers and luminescence imaging. To generate the first animal model, we subcutaneously implanted Nu/Nu mice with bioluminescent HCC827 human lung cancer cells that possessed a sensitivity to erlotinib (*22*). Erlotinib is an orally dosed small molecule drug that targets the epidermal growth factor receptor; its pharmacokinetics and pharmacodynamics occur on the timescale of hours (*22*–*24*). During the pre-treatment and treatment periods we compared the ability for our sensor, a caliper, and a luminescence imaging system to read out application readout device. Tegaderm and tissue glue were used to fix the sensor, battery, and holder on the mice. In our studies, we demonstrated that this wrapping protocol holds the sensors in place on the mice for at least one week. Eight days after tumor inoculation, when the tumor volumes were approximately 100 mm^3^, our sensor detected tumor growth over a 12-hour period by reading out an increase in resistance by a range of +21 to +64 ohms, with an average increase of 4.3 ± 2.2 ohms/hour (Mean ± SD) (n=6) (Figure 2a). During the tumor growth phase and before treatment, we ran an experiment characterizing the relationship between the sensor, the caliper, and the luminescence imager by ranking the readouts of each device according to readout magnitude (see Supplementary Fig. S4). After ranking measurement magnitudes three times over a seven-day period, the sensor and caliper measurements showed the closest correlation with an average rank difference of 1.59. The sensor and luminescence imager recorded an average rank difference of 1.74. Finally, the caliper and luminescence imager exhibited an average rank difference of 1.77.

**Figure 2:**
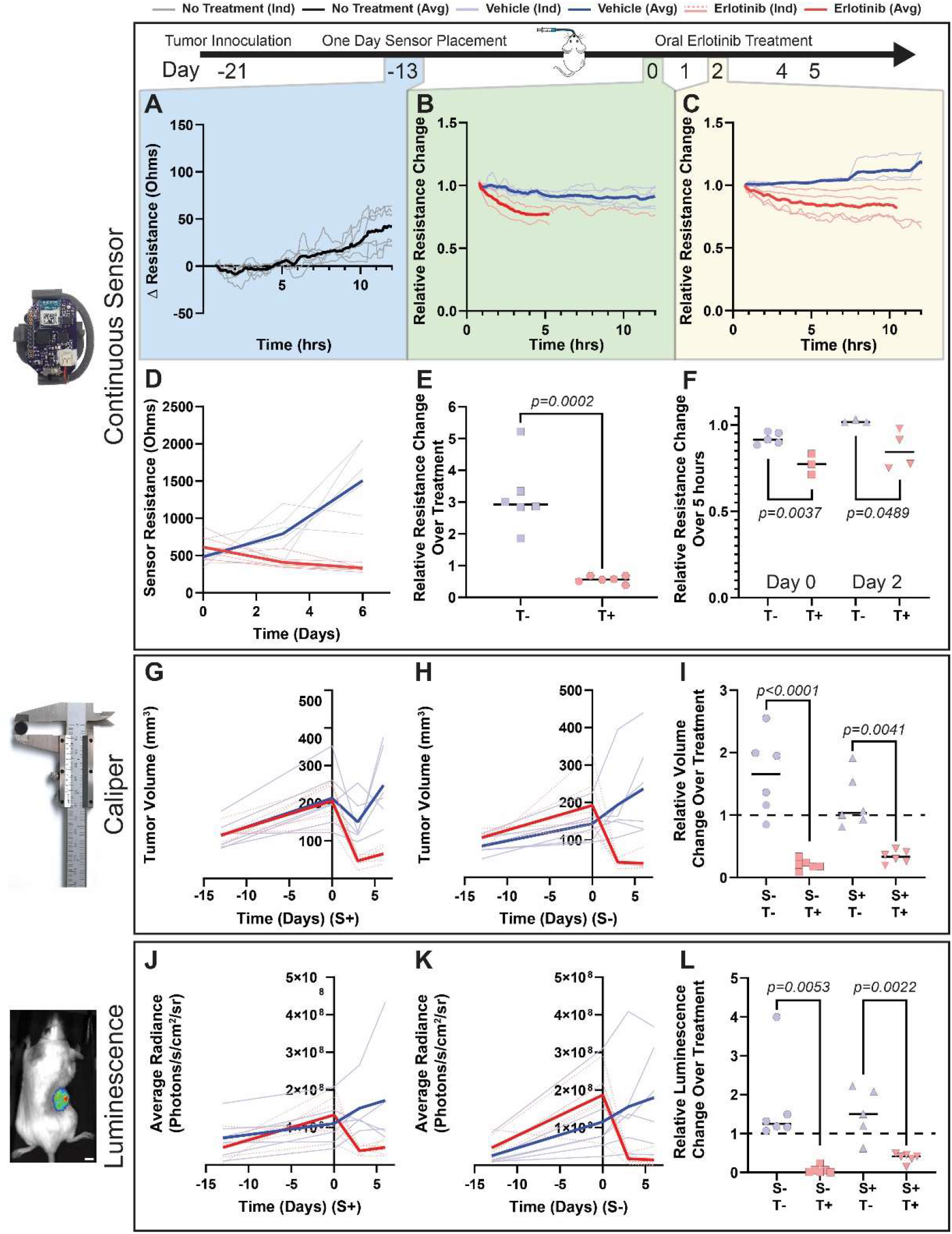
FAST sensor detects a decrease in tumor volume sooner than existing methods in HCC827 mouse models treated orally with erlotinib. (A-C) FAST reads out tumor volume progression continuously at 5 minute intervals in (A) Nu/Nu mice with ^~^100 mm^3^ subcutaneous HCC827 human lung cancer tumors receiving no treatment and (B,C) mice with ^~^200 mm^3^ tumors receiving 50 mg/kg erlotinib or vehicle treatments at intervals described in the figure. Individual mouse sensor trendlines are presented as 7 point moving averages. (D) FAST sensor measurements over entire treatment period. (E-F) Erlotinib and vehicle treated mice demonstrate significantly different sensor readouts over (E) the entire treatment period and (F) just five hours after treatment administration. (G-I) Calliper and (J-L) luminescence imaging confirm the tumor volume measurements recorded by FAST and demonstrate that wearing the FAST device does not affect the outcomes of the treatment experiments. (S+ = With FAST Sensor; S-= No FAST Sensor; T+ = Erlotinib Treatment; T-= Vehicle Treatment. Data is presented as individual datapoint or curves. Bold = Average.) (Unpaired Two-Tailed Student’s t-tests) (SB = 5 mm).

To evaluate the ability for FAST to measure biologically significant changes in tumor volumes in vivo during erlotinib treatment, we performed experiments controlling for the pharmacodynamic effects of the treatment and the mechanical effects of the sensor backpack. This required separating the mice into four groups to control for both the sensor and the treatment, and 6 mice were assigned to each group. FAST measurements, caliper measurements, and luminescence imaging measurements conveyed tumor shrinkage in all erlotinib treated mice throughout the six-day treatment period. These same measurement techniques also reported tumor growth in vehicle treated mice throughout the same time period (Fig 2). These trends were recorded irrespective of the presence of the FAST sensor. The FAST sensor, however, began detecting a change in tumor progression almost immediately following therapy administration, compared to the other measurement techniques which required several days to discern a biologically significant difference. Within five hours of placing the sensors on the mice, all vehicle treated mice demonstrated larger relative sensor readouts compared to the erlotinib treated mice (Fig 2b); this readout occurred again on a following dosage day as well (Fig 2c).

During this treatment session, we analyzed the impact of the mechanical stress placed on the sensor from the animal’s movement, and we assessed the impact of the mechanical stress placed on the tumor by the sensor. In this experiment, we utilized the 28 μm thick sensors presented in this paper; however, we found that during our testing several of the sensors lost their electrical connection, likely due to kinetic friction causing the gold layer to shed from the SEBS. Only sensors that recorded data are presented in figure 2, and no other data was removed from the analysis. By the end of the treatment, neither the caliper measurements nor the bioluminescence imaging recorded a significant difference between mice with or without the sensor (Fig 2g-l). However, in a separate experiment we noticed that the sensor began to constrain tumor growth if the tumors grew to the point where they reached the edges of the 3D printed housing. We also performed an analysis of the normal pressure exerted by the elastic sensor on the tumor, and we present this analysis in the supplementary information.

Histological evidence supports the rapid sensor classification of responsive and nonresponsive tumors using FAST by demonstrating that the tumors undergo modifications at the cellular level within hours after treatment administration (Figure 3). We compared histology samples from tumors undergoing the full erlotinib and vehicle treatment schedule with tumors excised five hours after erlotinib treatment initiation. Immunohistochemistry from tumors excised at the five-hour timepoint showed an upregulation of cleaved caspase-3, a marker for cell death. These same tumors also exhibited a downregulation in Ki67, of phosphorylated epidermal growth factor receptor, which is a direct pharmacodynamic response to erlotinib. Hematoxylin and Eosin stained histology from tumors undergoing the entire treatment schedule showed that erlotinib reduced the cell density in the tumor compared to vehicle treated tumors. In addition to the immunohistochemistry performed in this study, previous studies examining the pharmacokinetics and pharmacodynamics of erlotinib demonstrate that biological effects from the drug begin occurring within 5 hours in humans, in mice, and in cell culture (*22*–*24*). Moreover, little difference is seen between tumors that underwent the continuous sensor readout protocol compared to tumors where the sensor readout protocol was not administered. Hematoxylin and Eosin stained histology of skin where the sensors were placed for one week showed no signs of tissue damage.

**Figure 3:**
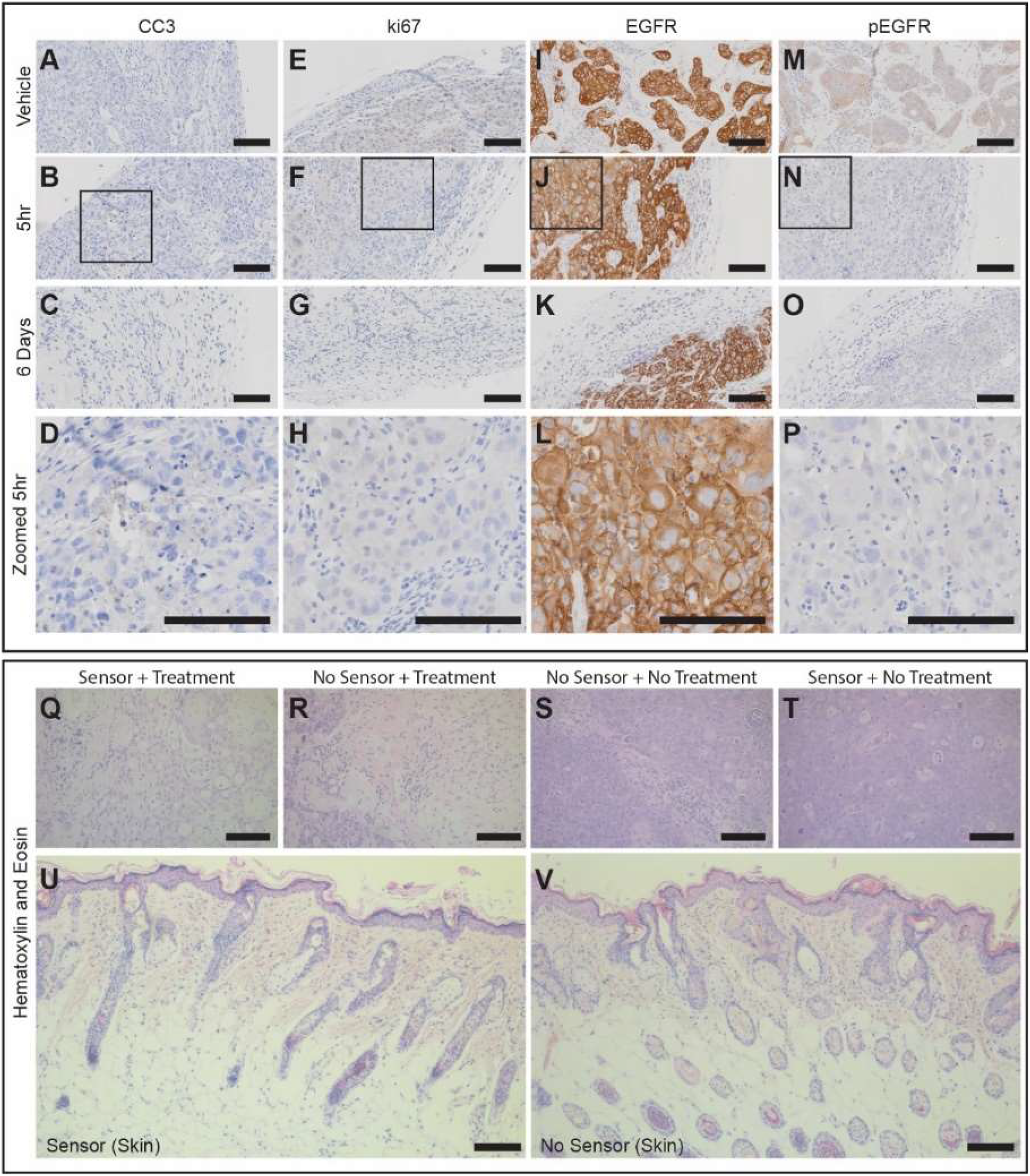
Histology from HCC827 tumors treated with erlotinib validates rapid FAST sensor readouts of tumor volume progression. (A-P) Immunohistochemistry of tumors excised from mice treated for 6 days with vehicle (Vehicle), treated for 5 hours with erlotinib, or treated for 6 days with erlotinib. Stains are for: Cleaved Caspase 3 (CC3), a marker associated with cell death; ki67, a marker associated with cell proliferation; Epidermal Growth Factor Receptor (EGFR); and Phosphorylated Epidermal Growth Factor Receptor (pEGFR). Erlotinib is an active inhibitor of EGFR and prevents phosphorylation. (Q-V) Hematoxylin and Eosin (H&E) stains of (Q-R) tumors and (U,V) skin from mice that did or did not wear FAST for 6 days. (SB = 100 μm).

In addition to characterizing the sensor with a clinically approved small molecule treatment, we also performed sensor characterization on an A20 B-cell lymphoma solid tumor model in Balb/c mice using an experimental immunotherapy. Specifically we treated the mice with an unmethylated CG–enriched oligodeoxynucleotide (CpG)-a Toll-like receptor 9 (TLR9) ligand-and anti-OX40 antibody via intratumoral injections (*25*). The sensor measurements in this tumor model were only directly compared to caliper measurements, because the presence of luminescence proteins in the cells generated an immune response that confounded the effects of the treatment. Similar to the last experimental model, the sensor was able to detect a change in tumor progression between treated and vehicle treated tumors within five hours after sensor placement. All immunotherapy treated tumors possessed a lower relative sensor readout than the vehicle treated tumors (Figure 4a,b). Additionally, vehicle treated tumors demonstrated a statistically significantly higher relative sensor readout compared to a control group where sensors were placed on animals without any tumors, while CpG and aOX40 tumors demonstrated a statistically significantly lower relative sensor readout compared to the same control group (Supplementary Figure S5). This demonstrates that mouse movements and breaths do not impact the ability for FAST to discern growth or shrinkage in in vivo tumors. Three weeks following therapy administration every treated tumor was completely eradicated, comparable to the results published previously on this therapy and tumor model (25). In this model, we utilized sensors with a 41 μm thick layer of SEBS, and all sensors performed unceasingly over the entire period of interest. Both the sensor and the caliper read out significant tumor shrinkage in immunotherapy treated tumors compared to vehicle treated tumors over the entire treatment period (Figure 4c-f).

**Figure 4:**
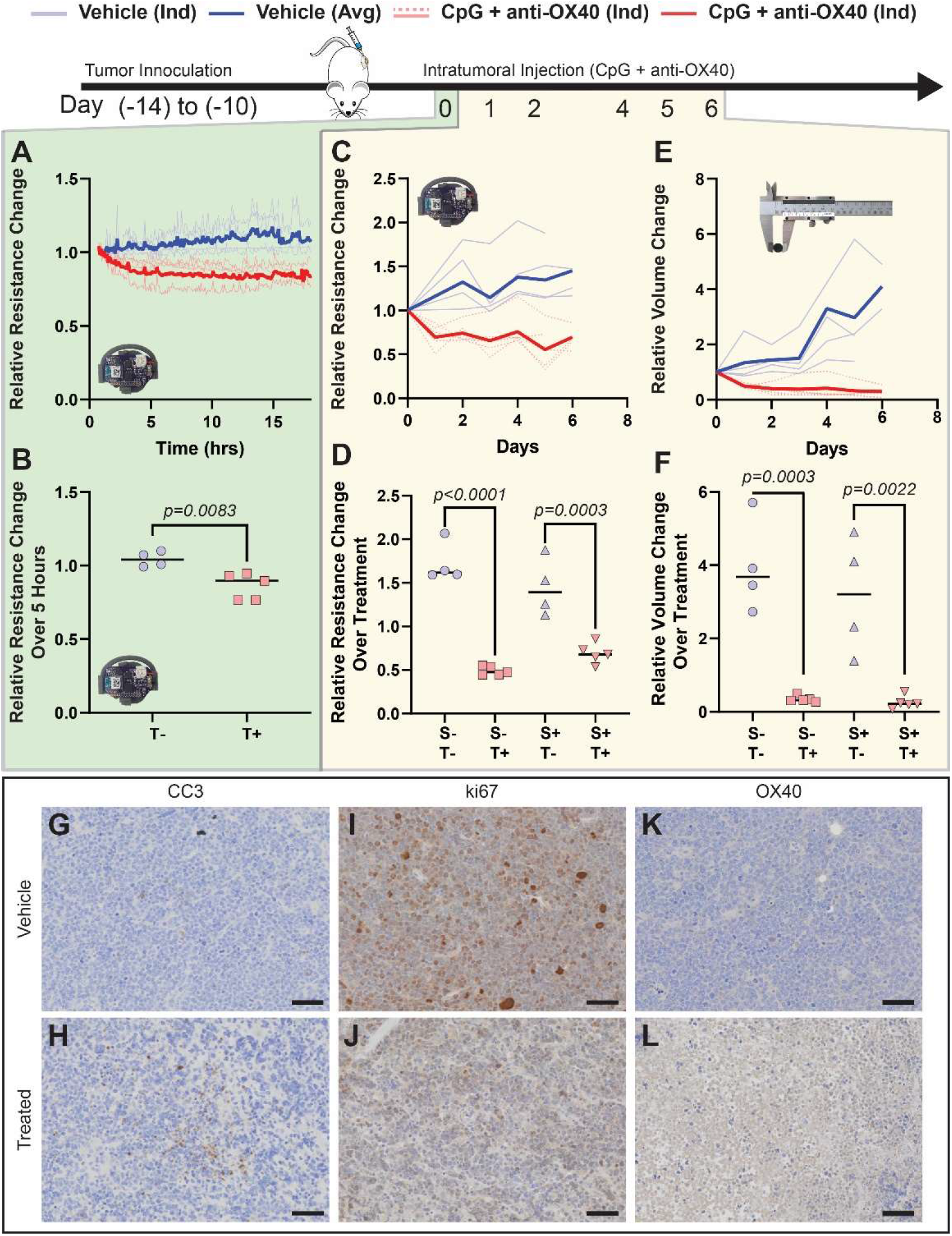
FAST sensor detects a decrease in tumor volume sooner than existing methods in A20 mouse models treated intratumorally with CpG + anti-OX40. (A-D) FAST reads out tumor volume progression continuously at 5-minute intervals in Balb/c mice with subcutaneous A20 B-cell lymphoma tumors receiving 40 μg of CpG and 4 μg of anti-OX40 (n=5) or vehicle (n=4) treatments over (A,B) the first few hours following treatment or (C,D) the entire treatment period. Individual mouse sensor trendlines are presented as 7 point moving averages. (E-F) Tumor volume measurements using callipers confirm FAST readouts over the entire treatment period. (G-L) Immunohistochemistry of tumors excised from mice treated for 6 days with vehicle (Vehicle) or treated once with CpG + aOX40 (Treated). Staining is against (G,H) Cleaved Caspase 3 (CC3), (I,J) ki67, and (K,L) OX40. The treated stains are from tumors excised within 6 hours after treatment initiation. (T+ = CpG and anti-OX40 Treatment; T-= Vehicle Treatment; S+ = With FAST Sensor; S-= No FAST Sensor. Scale bar = 50 μm. Data is presented as individual datapoint or curves. Bold Line = Average; B: Unpaired Two-Tailed Student’s t-test; D,F: One way ANOVA with Tukey’s multiple comparisons test.).

Immunohistochemistry from the A20 tumors demonstrates an immediate pharmacodynamic response following treatment initiation that supports the FAST measurement readouts. Specifically, the cell death marker Cleaved Caspase 3 was upregulated in tumors within 6 hours after treatment initiation (Figure 4g-h), but the cell proliferation marker ki67 was still present in the treated tumors at the 6-hour excision time point (Figure 4i-j). CpG also initiated an upregulation of OX40 within 6 hours after initiation, providing a target for the dosed antibodies to bind to and stimulate an immune response (Figure 4k-l).

## Discussion

In this paper, we presented a sensor system capable of continuously and accurately measuring solid tumor size progression, and we demonstrated its ability to discern treatment efficacy within just five hours after therapy initiation in two preclinical subcutaneous tumor models. Currently, experimental computational models are in development that predict a patient’s response to a given therapy (*26*–*28*), and combining these modeling tools with continuous tumor volume sensors may increase these algorithms’ sensitivities and specificities to clinically relevant levels. Moreover, next generation sequencing has enabled healthcare professionals to prescribe treatments that target specific cancer mutations, and clinical trials utilizing next generation sequencing as a screening tool have demonstrated objective responses in up to 20-40% of patients (*29, 30*). These trial outcomes highlight both the power and the limitations of targeted therapy screening and suggest that monitoring technologies such as FAST can provide information during treatment that cannot be assessed through sequencing alone. Importantly, our sensor focuses on measuring short-term primary tumor progression rather than metastatic progression, which provides a dataset that rapidly categorizes ineffective treatments by accurately capturing primary tumor growth. For potentially effective treatments in which FAST sensors rapidly read out a reduction in the primary tumor’s volume, the sensor data can be utilized as an indicator to perform follow-up screenings that provide additional information on tumor progression that confirm a reduction in total tumor burden. Of note, some tumors are known to undergo pseudoprogression after treatment initiation, a phenomenon where the tumor grows for a period of time preceding subsequent regression, and the occurrence of tumor growth does not necessarily signify a failed therapy (*31, 32*). In our studies, we directly compared the tumor progression of vehicle and drug treated mice, providing appropriate controls to ensure confidence in our measurements. While our sensors did not detect tumor pseudoprogression during the treatments we studied in this paper, future work may enable us to detect differences between normal progression and pseudoprogression growth rates using the real-time data generated by our sensor.

Of note, while we developed an encapsulated version of the sensor that can withstand contact with fluid (Supplementary Figure S6), the size limitations of a mouse model prevent the implantation of FAST. This limited our experiments to testing on subcutaneous tumors. Efforts to translate the sensor to humans should consider the surgical impact associated with placing the sensor at a given tumor location, as the sensor may best be suited for easily accessible tumors. Further work optimizing the battery life and size of the associated electronic printed circuit board is also required in pursuit of a fully implantable system. Passive wireless sensing systems may provide an alternative path to the implementation of implantable systems (*33, 34*). Still, this sensor’s ability to continuously, autonomously, and accurately record tumor volume progression suggests that this method could supplant current tumor progression measurement techniques, unlocking new avenues for drug discovery screenings, clinical cancer therapy assessments, and basic cancer research that take advantage of the sensor’s in vivo time dependent datasets.

## Acknowledgements

We thank members of the Bao, Gambhir and Hiesinger labs for helpful discussions on the project. We thank Dr. Israt Alam for advice about cell culture and the A20 tumor model. We thank Boris Murmann and Nicholas Vitale for their contributions to the design of the printed circuit board. We acknowledge the Stanford Center for Innovations in In vivo Imaging (SCi^3^) - small animal imaging center and the Stanford Animal Histology Services for supporting the imaging and histology performed in this article. We acknowledge the Stanford Veterinary Services and Monika Huss for help designing the animal experiments. Part of this work was performed at the Stanford Nano Shared Facilities, supported by the National Science Foundation under award ECCS-1542152. The photo of the hand with the cellphone in figure 1 is provided with the permission of Facebook Design Resources. The photo of the calipers in figures 2 and 4 was distributed by WikiCommons under a Creative Commons 3.0 license and was taken by Simon Eugster.

## Author Contributions

A.A., P.M., S.S.G. and Z.B. designed the project and the experiments. A.A. and N.M. designed the strain sensor. A.A. designed the sensor backpack device. A.A. and A.M.B. performed in vitro characterization of the sensors. A.A. and Y.K. designed the printed circuit board and cell phone app. C.T.C., R.F. and R.S. performed cell culture. A.A. and C.T.C. performed the animal experiments. A.A., J.D., and Z.B. wrote the manuscript. All authors reviewed and commented on the manuscript.

## Data Availability

The authors declare that the data supporting the findings of this study are available within the paper and its supplementary information files.

## Code Availability

Custom code used to program the custom designed printed circuit board is available from the corresponding author upon request.

## Materials and Methods

### Sensor Backpack Fabrication

A schematic of the sensor and its fabrication process is located in Supplementary Figure S1. Sensors were fabricated on a 5.0 cm × 7.5 cm glass slide (Fisher Scientific, Waltham, USA). As an anti-stick coating, a Micro-90 solution (Cole-Parmer, Vernon Hills, USA) was coated on a slide by spin coating 300 μL of solution on the slide at 600 rpm for 20 seconds. A WS-650MZ-23NPP spin-coater from Laurell Technologies (North Wales, USA) was used. Solutions of 33 mg/mL and 50 mg/mL SEBS (Asahi Kasei, 1221, Chiyoda City, Japan) in Cyclohexane (Fisher Scientific) were generated, and the solution was mixed overnight. The SEBS solution was then drop casted on a 3 inch × 2 inch glass slide. To create the 28 μm thick substrate, 4 mL of 33mg/mL solution was used. To create the 41 μm thick substrate, 4 mL of 50mg/mL solution was used. To create the 72 μm thick substrate, 4 mL of 50mg/mL solution and 2 mL of 33mg/mL solution were combined and used. A transparency film (Acco Brands, Boonville, USA) mask was mechanically cut using a Cricut machine (South Jordan, USA) from a mask designed in Solidworks (Dassault Systemes, Velizy-Villacoublay, France). The sensor design consisted of an 11 mm × 1.5 mm strip, book ended by 3 mm × 3 mm connection pads. Once cut, the transparency film was sprayed with a non-stick Teflon spray (Dupont, Eleutherian Mills, USA) and placed on the SEBS substrate. Then, a 50 nm layer of gold was deposited on the SEBS at 0.6 angstroms/second using a metal evaporator from Thermonionics Laboratories Inc (Hayward, USA). Gallium-Indium eutectic (Sigma Aldrich, St. Louis, USA) was placed on the connection pads and a 30 G multicore wire (McMaster Carr, Elmhurst, USA) was attached to the connection pad using paper tape. The wires were then soldered to a custom designed printed circuit board (See Supplementary Figure S3) assembled by Digicom Electronics (Oakland, USA). The circuit board is powered by a 150 mAh Lithium-Ion rechargeable battery (Digikey, Thief River Falls, USA). When awake, the average current draw for the circuit board is 3.5 mA. The sensor backpack (See Supplementary Figure S1) was printed in 3 pieces on a Formlabs Form 2 Printer (Sommerville, USA). The two rigid rods were printed in either Rigid resin or Grey resin, while the flexible base was printed in Flexible resin.

Fully encapsulated sensors were fabricated by first spin coating a polydimethylsiloxane (PDMS) (Sylgard 184, Dow, Midland, USA) layer mixed at a 10:1 ratio (PDMS : crosslinker) at 1000 rpm for 30 seconds. The PDMS was then cured at 70°C for 12 hours. Then a 40 nm thick gold film was evaporated onto the PDMS substrate at 0.5 A/s. This gold film was sandwiched between two 3 nm thick evaporated chromium films and patterned using the transparency shadow masks described above. Gallium-Indium eutectic (Sigma Aldrich) was placed on the sensor connection pads along with a 36 G multicore wire (McMaster Carr). The entire device was then fully encapsulated in Kwik Sil (World Precision Instruments, Sarasota, USA). We demonstrated that the device could remain in contact with Phosphate Buffered Saline and with mouse tissue in euthanized mice while maintaining its conductivity and ability to read out strain measurements through changes in resistance between 0% to 40% strain.

While stretchable sensors are known to undergo hysteresis and experience drift during repeated cycling, the fact that this application of the sensor only requires one stretching cycle eliminates the potential for error associated with these materials-based concerns. Moreover, the viscoelastic properties of SEBS causes the sensor to experience a reduction in resistance over time (Supplementary Figure S2b), but the sensor nears equilibrium approximately 30 minutes after strain is applied. For this reason, in vivo measurements were normalized to the data points taken 30 minutes or more after sensor placement. Placing the sensors on a 3D object compared to providing strain in one dimension may affect the exact readouts of the sensor; however, the data in figure 1g demonstrates that an increase in resistance is still exponentially proportional to an increase in the ellipsoid shape that the sensor is wrapped around. Finally, animal movement does cause the sensor to constantly undergo small changes in strain; however, these small changes in strain are averaged out over multiple points and have been shown through our measurements to not affect the statistical significance of the in vivo experiments (Supplementary Figure S5).

### Sensor in vitro Characterization

To measure the resistance during stretching, we attached samples to a homemade stretching station and connected the samples to an LCR meter (Keysight Technologies, E4980, Santa Rosa, USA). Before beginning the measurements, sensors were stretched to 200% strain by hand more than 20 times. Samples were then stretched between 0% and 100% strain at 1% intervals, approximately 120 μm per step, and resistance measurements were recorded in LabView (National Instruments, Austin, USA). Following this test, the samples were then stretched to 50% strain, and the resistance of the sensor was measured over the course of 45 minutes. This test demonstrated that although the sensor underwent relaxation over time, much of the relaxation occurred within the first 45 minutes (see Supplementary Figure S1). After this test, the sensor was then stretched from 50% to 60% strain at 0.083% intervals, approximately 10 μm per step.

To measure the force required to strain the sensor to a given length, we attached the samples to an Instron 5565 (Norwood, USA). We stretched the samples at a rate of 50 mm/min, zeroing the displacement and the force once the sample reached 0.05 N of force. Forces were recorded using a 100 N force gauge provided by Instron and read out on the machine’s accompanying software. Each sample was stretched until its breaking point.

To measure the thickness of each sensor, we used a Bruker Dektak XT-A profilometer (Billerica, USA) and took the average of 10 different reading from multiple sensors taken from various locations on the sensor. The edges of the sensor tended to have a slightly thicker measurement compared to the center of the sensor, leading to a slight variability in thickness readouts (see Supplementary Figure S1).

To measure the ability of FAST to read out the variation in volume of different shapes, we 3D printed ellipsoid shapes cut in half down their center line. All shapes were scaled linearly and possessed heights between 2.5 mm and 5.6 mm, as measured using calipers. These shapes were designed in Solidwork and printed on an Ultimaker 3 using Ultimaker PLA filament (Geldermalsen, Netherlands). The FAST devices were placed on the shapes, and the sensors were allowed to relax for 20 seconds before the resistance measurement was recorded.

### Subcutaneous HCC827 tumor treatment with Erlotinib

All animal procedures were approved by the Stanford Institutional Animal Care and Use Committee and conducted in accordance with Stanford University animal facility guidelines. The HCC827 human lung cancer cell line was obtained from ATCC (CRL-2868, Manassas, USA) and was then transfected with the firefly luciferase reporter gene. Before injecting the cells into mice, the cells were tested and shown to be pathogen free by the Stanford Department of Comparative Medicine Veterinary Service Center (Stanford, USA). Five million cells were injected into the right flank of six- to eight-week-old Nu/Nu mice (Charles River Laboratories, Wilmington, USA) after being mixed with Matrigel (Corning, Corning, USA). Mice were housed in the Laboratory Animal Facility of the Stanford University Medical Center (Stanford, CA).

The sensors were placed on six of the animals once the tumors reached a size of approximately 100 mm^3^ and were left on the animals for one day. When placing the sensors on the animals, the mice were anesthetized with 1-3% isoflurane. Buprenorphine Sustained release was also dosed to the animals at 0.5-1.0 mg/kg. Before beginning the procedure, we checked the absence of paw reflexes by pinching a hind paw with tweezers and checked the absence of eye reflexes to make sure that the animal was fully anesthetized. A protective eye liquid gel (GenTeal, Alcon, Geneva, Switzerland) was then applied to the eyes with a cotton-tipped swab. If necessary, we then shaved the location where the sensor was to be attached to the animal around the tumor. The skin was then aseptically prepared with alternating cycles of betadine or similar scrub and 70% ethyl alcohol. Using a surgical tissue glue (3M, Saint Paul, USA) the sensor was attached to the skin of the animal so that the tumor was positioned in the center of the sensor. A 1.3 inch in diameter tegaderm wrap was then applied on top of the sensor and to the animal’s skin so that the sensor remained snuggly attached to the animal. The battery was similarly attached to the skin using tegaderm and was placed on the opposite flank of the sensor. Every day that the sensor remained on the animal the battery was replaced and the tegaderm wrap was replaced above the battery.

Once the tumor reached a volume of approximately 200 mm^3^, the mice were broken up into four groups of six: one group received the erlotinib treatment and the sensor protocol; one group received the erlotinib treatment and did not receive the sensor protocol; one group received a vehicle treatment and did receive the sensor protocol; and one group received a vehicle treatment and did not receive the senor protocol. The treated mice were dosed with erlotinib hydrochloride (Fisher Scientific) dissolved in a mixture of Captisol (Selleckchem, Houston, USA) and water. Erlotinib was dosed at 50 mg/kg via an oral gavage to mice. Mice that did not receive the erlotinib were dosed with vehicle only. Dosing occurred on days 0, 1, 2, 4 and 5. On day 3, mice did not receive treatment and they also did not receive the sensor protocol. Diet gel 76A and sterile water gel (ClearH20, Westbrook, USA) were placed in the mouse cages to ensure easy access to food and hydration. The weight of each mouse was recorded over time and this data is presented in Supplementary Figure S7. Mice wearing the sensor were singly housed to prevent other mice from chewing through the sensor backpack. On days 0, 3, and 6, all mice underwent caliper measurements (McMaster Carr), individual time-point sensor measurement, and bioluminescence imaging. Luminescence imaging was performed on a Lago X (Spectral Instruments Imaging, Tucson, USA), and image analysis was performed in the accompanying Aura software. The mice were euthanized on day 6, and the tumors and skin next to the sensors were harvested for histology.

The excised tissues were fixed in a 4 % Paraformaldehyde (PFA) solution for 24+h, followed by 70% ethanol for 24+h. Immunohistochemistry staining utilized the following antibodies: EGFR (D38B1) XP Rabbit mAb #4267 (Cell Signaling Technology, Danvers, USA); Phospho-EGF Receptor (Tyr1068) (D7A5) XP Rabbit mAB #3777 (Cell Signaling Technology); #9579 Cleaved Caspase-3 (Asp175) (D3E9) Rabbit mAB (Cell Signaling Technology); ki67 Polyclonal antibody #27309-1-AP (ProteinTech Group, Rosemont, USA); Biotinylated Goat Anti-Rabbit IgG (H+L) (ab64256) (Abcam, Cambridge, United Kingdom). HRP-Conjugated Streptavidin was purchased from ThermoFisher. DAB Substrate Kit ab64238 was purchased from Abcam. Antigen retrieval was performed by incubation for 20 minutes in pH 6.0 citric acid at 100°C. Antibody dilutions and staining procedures were performed as suggested by the manufacturer.

### Subcutaneous A20 tumor treatment with CpG and anti-OX40

All animal procedures were approved by the Stanford Institutional Animal Care and Use Committee and conducted in accordance with Stanford University animal facility guidelines. The A20 B-Cell lymphoma cell line was obtained from ATCC (TIB-208). Before injecting the cells into mice, the cells were tested and shown to be pathogen free by the Stanford Department of Comparative Medicine Veterinary Service Center (Stanford, USA). Five million cells were injected into the right flank of six- to eight-week-old Balb/c mice (Charles River Laboratories, Wilmington, USA). Mice were housed in the Laboratory Animal Facility of the Stanford University Medical Center (Stanford, CA). As described in the HCC827 methods section, mice were split into treatment and vehicle groups, and the sensors were applied to all the animals. Caliper measurements and sensor measurements were recorded daily over the span of 6 days. On days 0, 1, 2, 4, 5, and 6, the treated animals were injected with 40 μg of CpG ODN 2395 (Invivogen, San Diego, USA), a class C tlr9 ligand and 4 μg Anti-OX40 (CD134) monoclonal antibody (rat immunoglobulin G1 (IgG1), clone OX86) (BioXCell, Lebanon, USA). The total volume injected in the treated and vehicle treated mice was ^~^13-16 uL and varied depending on the concentration of the antibody. The weight of each mouse was recorded over time and this data is presented in Supplementary Figure S8. On day 3, the sensor was removed from the animal and no therapy was given to the animal. Supplementary Figure S8 also includes data showing the tumor progression and regression of drug treated and vehicle treated mice that did not continuously wear the FAST sensor. Diet gel 76A and sterile water gel (ClearH20, Westbrook, USA) were placed in the mouse cages to ensure easy access to food and hydration. Mice wearing the sensor were singly housed to prevent other mice from chewing through the sensor backpack.

### Tumor Compression Experiments

Using an Instron Machine, two steel compression platens compressed an excised tumor at 2 mm/min. Excised tumors were tested on the machine within 1 hour following euthanasia, and the tumors were kept in Phosphate Buffered Saline after excision and before testing. Both vehicle and drug treated A20 tumors were dosed via an intratumoral injection one day before tumor excision. The force versus displacement readouts were recorded on the accompanying Instron software and are presented in Supplementary Figure S9. Of note, the software began recording once the force gauge read out a value of at least 3 mN.

### Statistical Analysis

No data was excluded from the analysis. Paired and unpaired two tailed student’s t-tests and One-way Anova tests with Tuckey’s multiple comparisons tests were performed using Prism Version 8.3 (GraphPad) or Microsoft Excel (Microsoft). Paired t-tests were used when performing direct comparisons between individual sensors at different strains. Unpaired t-tests were used in other situations in which a paired t-test was not appropriate. A value of P < 0.05 was considered statistically significant. Figure captions and text describe the number of replicates used in each study. Figure captions define the center line and error bars present in the plots.

