## Supplementary Information for "A flexible electronic strain sensor for the real-time monitoring of tumor progression"

**Affiliations:**

^$^Deceased

**This PDF file includes:**

**Supplementary Text:** Estimating the pressure exerted by the FAST device on the tumor

**Supplementary Text:** Using the FAST device to measure ellipsoid volume

**Supplementary Fig. S1:** FAST fabrication design schematics.

**Supplementary Fig. S2:** Strain sensor characterization.

**Supplementary Fig. S3:** Circuit Board Design and testing.

**Supplementary Fig. S4**: Ranked comparisons of tumor volume measurement tools.

**Supplementary Fig. S5:** In vivo Sensor Measurement Controls.

**Supplementary Fig. S6**: Encapsulated PDMS Strain Sensor Characterization.

**Supplementary Fig. S7**: Mouse weight for HCC827 model over treatment period.

**Supplementary Fig. S8**: Mouse weight for A20 model over treatment period.

**Supplementary Fig. S9**: Estimates of force and pressure on tumor from sensor.

Planning for Device Scale-Up

Although the devices were manufactured in small batches using rapid prototyping tools, the device can be scaled up for manufacturing with only minor changes. Large uniform sheets of SEBS can be manufactured using extrusion. Gold evaporation and PCB manufacturing are already performed at large scales. The 3D printed rigid rods and flexible base can be injection molded with minor modifications to eliminate unmoldable geometries. When 3D printed, the long and short rigid rods require 1.61 mL and 0.94 mL of resin, including supports, and the flexible base requires 0.89 mL of resin. This means each part costs $0.24, $0.14, and $0.18 in resin, respectively, and would cost <$0.01 each when injection molded. Even in small batches, the PCB design costs $49 to manufacture, and this cost can be reduced significantly at higher volumes. The rechargeable battery retails for $4. At $60/g the 0.033 mg of gold used in a single sensor would cost <$0.01, and at $2/kg the 7 mg of SEBS in the sensor would also cost <$0.01. Therefore the majority of the cost lies in the re-usable PCB and battery rather than in the other sensor components, but it is possible to manufacture the device in a laboratory for <$60 each in raw material costs.

Estimating the pressure exerted by the FAST device on the tumor

Using the force versus strain curves in Figure 1, it is possible to estimate the force and pressure being applied to the tumor by the elastic sensor. The sensor is placed on the FAST backpack and pre-strained to a strain of 50%. The sensor undergoes relaxation following the pre-strain event, but we will use the values in Figure 1 so that we provide an estimate for the maximum force applied. The normal force on the tumor can be calculated using Equation 1, where F is the force applied to the tumor, T is the total tension on the elastic sensor, and θ is the acute angle generated between the elastic sensor and the mouse skin.

Equation 1: $F=2Tsin(\theta)$

Equation 2 is used to calculate the tension, where k is the spring constant of the elastic sensor at 50% strain, l is the total length of the pre-stretched sensor before it is applied to the tumor, b is the height of the tumor, and T_0_ is the initial tension on the sensor before it is stretched to fit the tumor.

Equation 2: $T=k\left( \frac{l}{\cos\left( \theta\right)}-l \right)+T_{0}$

Equation 3 is used to calculate the angle θ.

Equation 3: $\theta={tan}^{-1}(\frac{b}{l})$

To calculate the pressure, the cross-sectional area (A) of the tumor was evaluated using Equation 4, and the force was divided by the cross sectional area.

Equation 4: $A=\pi{(4b)}^{2}$

For the 28 µm thick SEBS sensor, k = 0.79 N/m and T_0_ = 0.064 N. For the 41 µm thick SEBS sensor k = 2.06 N/m and T_0_ = 0.185 N. For both sensor thicknesses l=0.018 m and b is variable between 0.002 to 0.006 m. This assumes that the height of the tumor is approximately 4x smaller than the length and width of the tumor.

Using these equations, an estimate of the Force and pressure exerted on the tumor was calculated and is presented in Supplementary Figure S9. Compression tests on drug treated and vehicle treated excised A20 tumors showed no statistically significant difference at the force values provided by the sensor. Moreover, no tissue rupture events were recorded by the compression tests at these force values. Along with our tumor volume progression data, this experiment suggests that the force provided by the strain sensor to the tumor has little effect on the dynamics of tumor volume progression.

Using the FAST device to measure ellipsoid volume

It is possible to calculate the volume of an ellipsoid using a set of three strain sensors. Our FAST circuit board supports the use of three continuous sensor readouts, and our FAST backpack enables three strain sensors to be placed around the tumor at a given time. The first two strain sensors are placed over the tumor so that they are perpendicular to each other. The third sensor is placed so that it wraps around the tumor parallel to the mouse’s skin. This third sensor is attached to the same rigid rod as another one of the sensors, but it is attached to the other side of the connection lip. Each of these three sensors reads out half of one of the three characteristic perimeters that make up the ellipsoid. Each of these perimeters (p_x_) is defined by two of the three characteristic radii (a,b,c) of the ellipsoid. The calculations for the volume (V) and perimeter of the ellipsoid are found in equations 5 and 6 below. The exact calculation for the perimeter of an ellipse is an infinite series, so instead we use a well-established approximation.

Equation 5: $V=\frac{4\pi}{3}abc$

Equation 6: $p\sim\pi\left[ 3\left( a+b \right)-\sqrt{(3a+b)(a+3b)} \right]$

It is possible for the sensor to wrap around the tumor in two different ways (See Supplementary Figure S9). In the first way, the sensor only contacts the highest portion of the tumor, like a tent and a tent pole. In the second way, a flexible band is placed around the tumor that grows and shrinks as the tumor volume progresses. This flexible band holds the sensor against the tumor so that it completely wraps around the tumor. In the first way, the height of the tumor can be calculated using Equation 7. It is not possible to calculate the perimeter of the characteristic ellipse that this sensor fitting is measuring. In the second way, the perimeter of the characteristic ellipse can be calculated using Equation 8. Using a standard curve for the strain sensor that characterizes how the change in strain affects the change in resistance (eg. Fig 1b), it is possible to calculate the sensor’s stretched length (l_1_) from the recorded resistance. The pre-stretch length (l) remains constant.

Equation 7: $l_{1}-l=2\sqrt{\left( \frac{l}{2} \right)^{2}+d^{2}}-l$

Equation 8: $l_{1}-l=\left( l-2a \right)+\frac{p}{2}$

In order to solve for the characteristic radii (a,b,c), the three simultaneous sensor readouts can be used to create a series of three equations, and these can be solved using an equation solver, such as the one provided in MATLAB. These three characteristic radii can then be plugged into Equation 5 to calculate the tumor volume.

Of note, the flexible band used to hug the sensor to the tumor can interfere with the gold coating on the SEBS elastomer and cause points of large strain. For this reason, the bands were not used during the in vivo experiments. However, just using the first wrapping method provided us with enough information to determine the difference in tumor volume progression between treated and vehicle treated tumors. The sensor resistance measurement itself was used in this paper as a stand-in for tumor volume, just as Radiance is used as a stand-in for tumor volume when performing luminescence imaging.

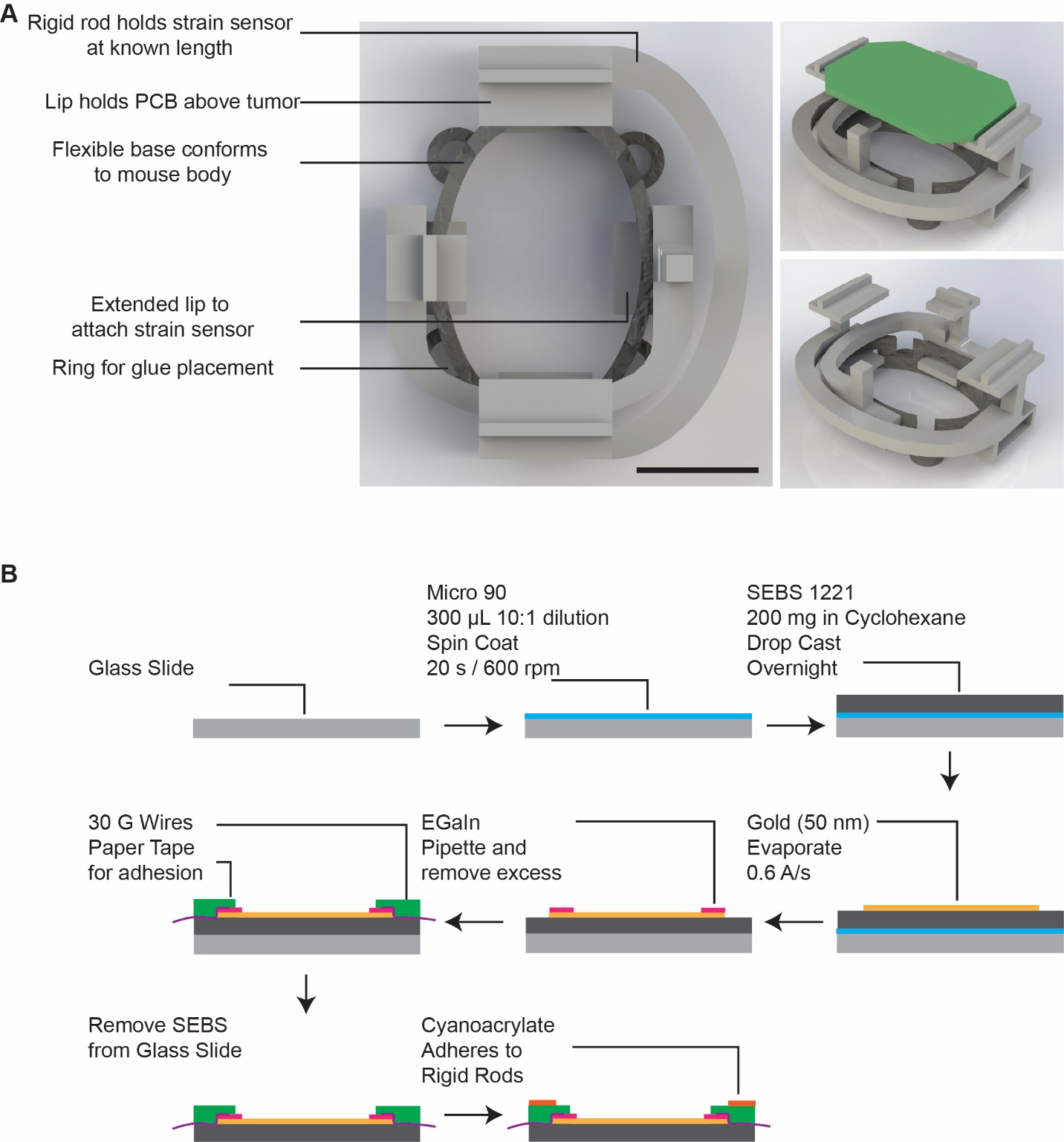

**Supplementary Figure S1: FAST fabrication design schematics.** (A) Solidworks rendering of FAST backpack. The green part in the top right image represents the printed circuit board. (SB = 10 mm). (B) Process used for fabricating cracked gold strain sensors.

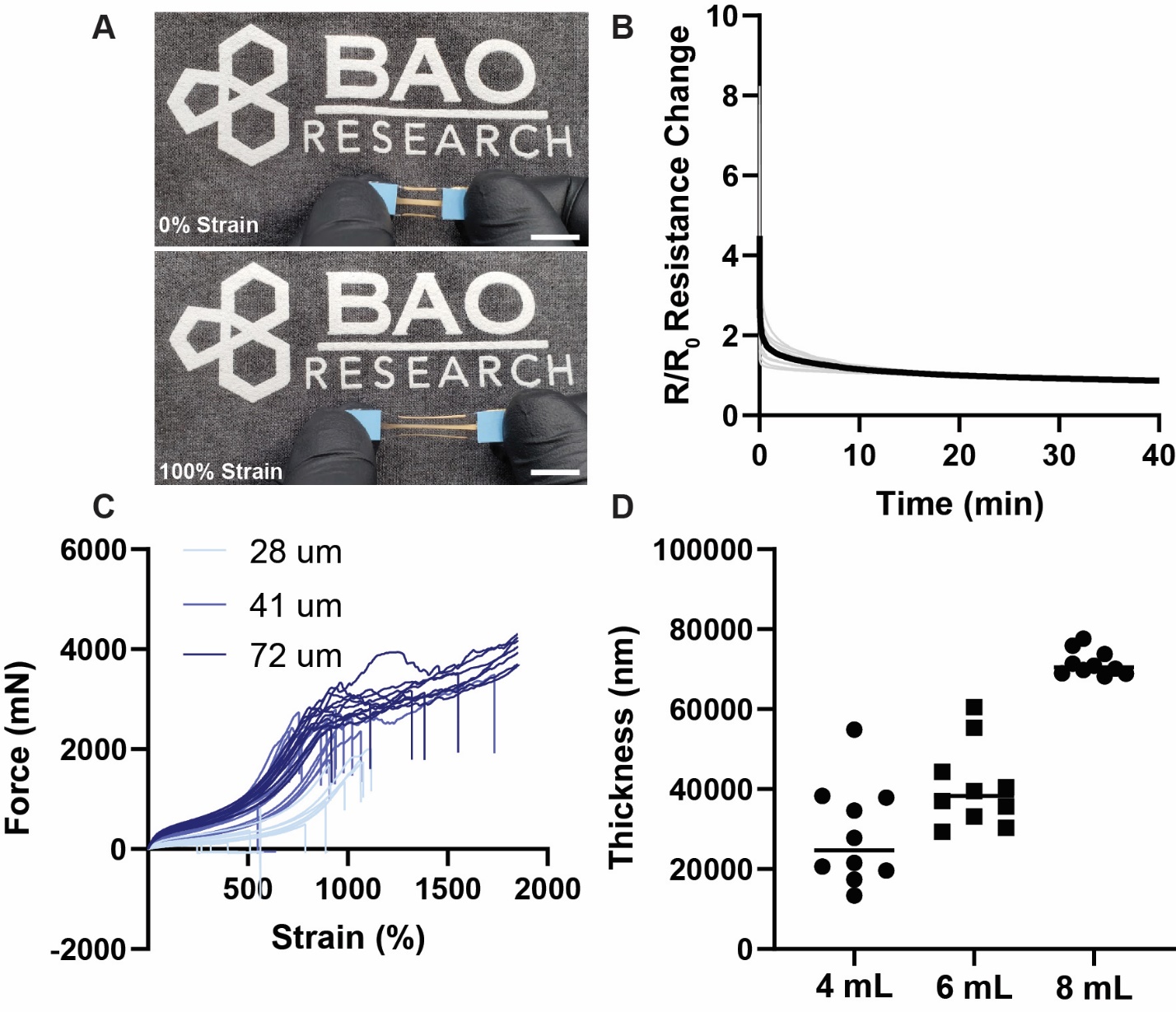

**Supplementary Figure S2: Strain sensor characterization.** (A) Optical image of cracked gold strain sensor being stretched to 100% strain (SB = 10 mm). (B) SEBS relaxation over time causes a decrease in resistance readings over time when stretched to 50% strain. This relaxion effect levels out at approximately 40 minutes (Individual curves, Bold Line = Average; n=14 sensors). The individual curves are standardized to the resistance reading at 20 minutes post strain event. (C) Instron tensile test of SEBS sensors with different thicknesses stretched until breaking point (n=12-13 sensors per group). (D) Profilometer measurements of SEBS sensor thicknesses. Sensors were fabricated with a varying amount of drop casted SEBS solution dissolved in cyclohexane (n=10 measurements).

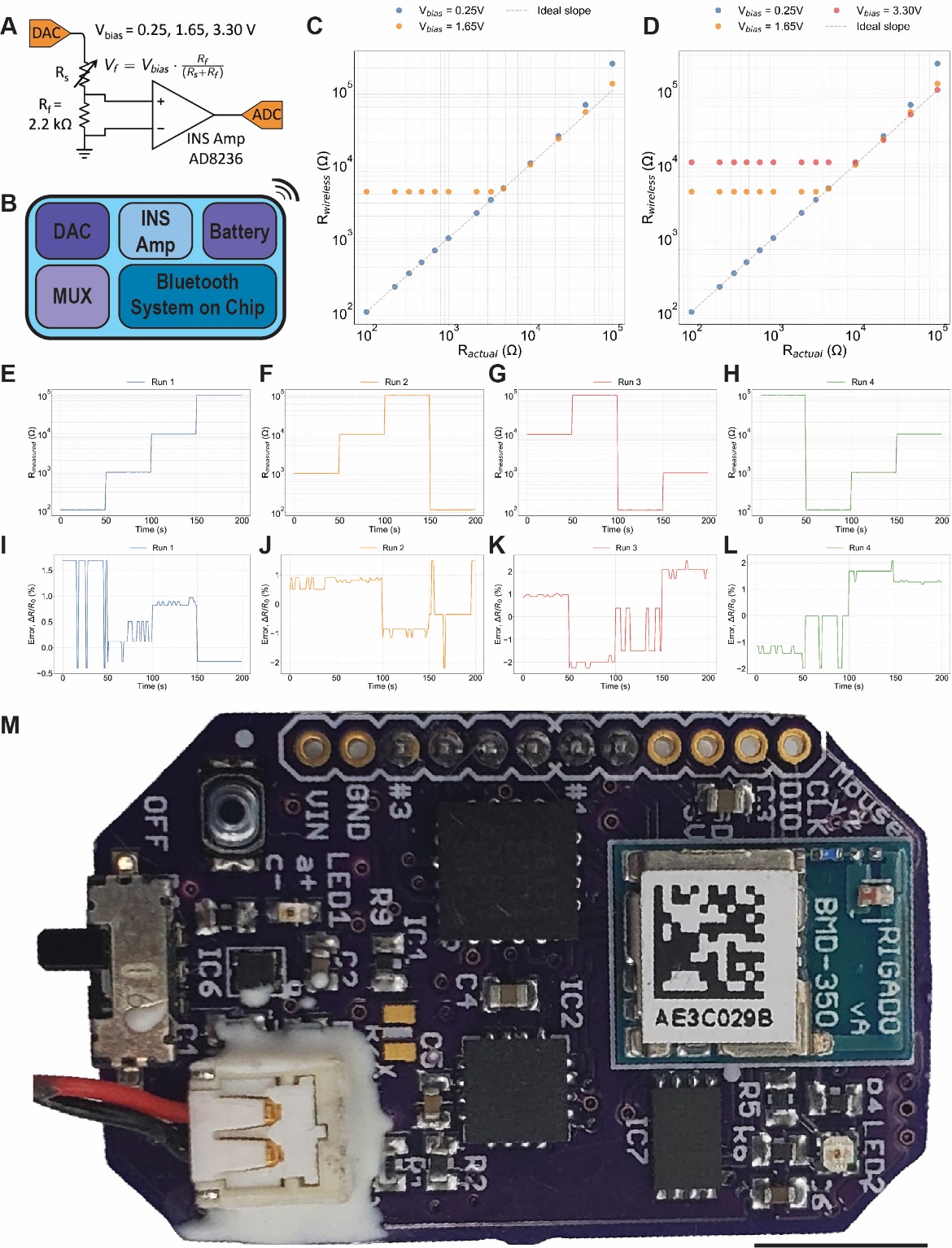

**Supplementary Figure S3: Circuit Board Design and testing.** (A) Sensor Readout diagram. A dynamic voltage range (V_Bias_) was used over the circuit containing the sensor resistor (R_S_) and a known resistor (R_f_). (B) The circuit board contains a digital-to-analog converter (DAC), a multiplexer (MUX), an Instrumentation Amplifier (INS Amp), a battery, and a Bluetooth system on a chip in order to record, read out, and transmit the sensor data. (C,D) A variable bias is passed through the circuit in “A” in order to reduce the error over the entire range of sensor resistances between 300Ω-60kΩ. (E-H) It is possible to switch among 3 channels with different resistances while (I-L) maintaining a readout error at or below 2%. (M) Photo of printed circuit board. SB = 5 mm.

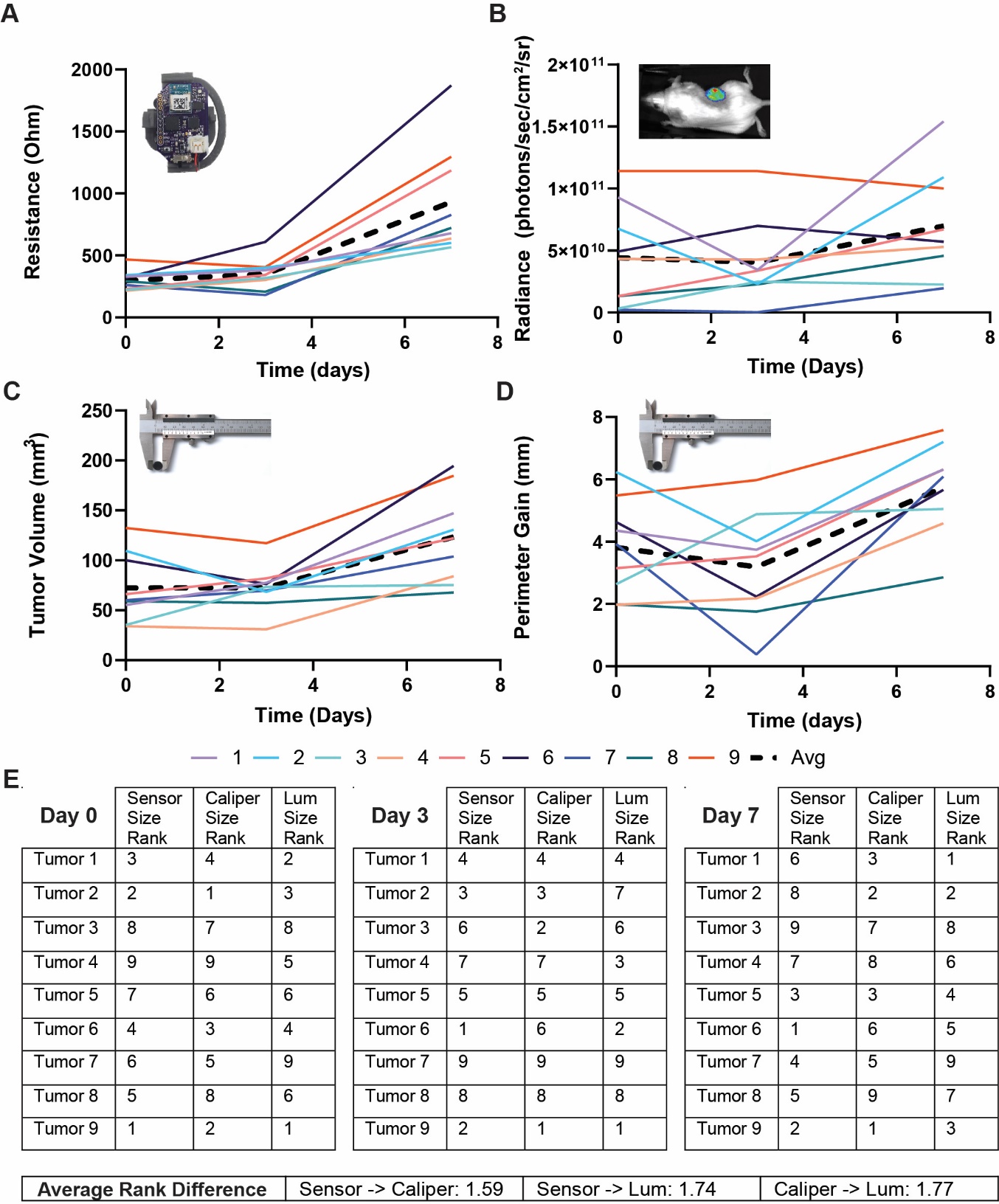

**Supplementary Figure S4: Ranked comparisons of tumor volume measurement tools.** (A) FAST resistance readouts, (B) Bioluminescence readouts, and (C) caliper tumor volume quantification readouts for growing subcutaneously implanted HCC827 tumors in Nu/Nu mice. (D) The calculated increase in the sensor length from a resting position when placed on a given tumor. See Supplementary information for methods on calculations. (E) The size rank of all the tumors on each day are tabulated based on each measurement technique and compared. The average rank difference for each of the methods over all the days is presented.

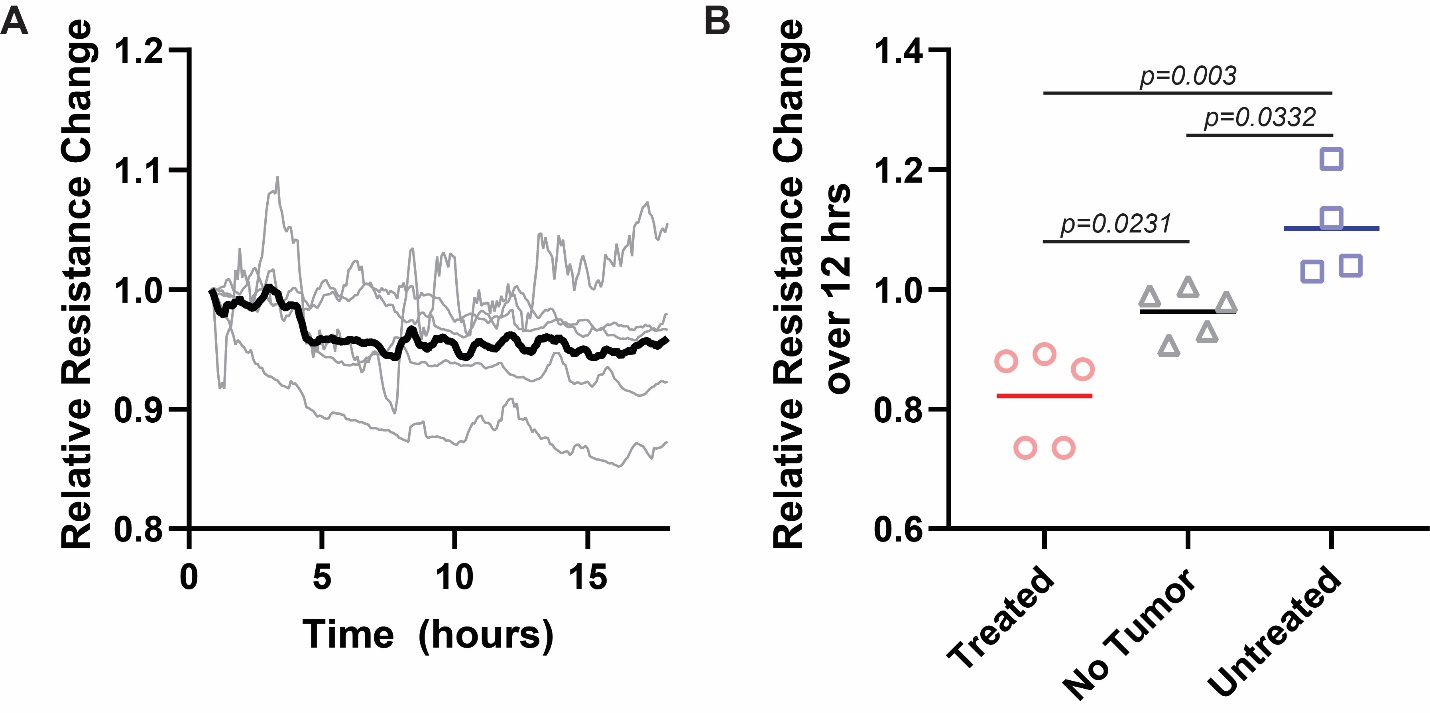

**Supplementary Figure S5: In vivo Sensor Measurement Controls.** (A) The relative resistance change over time for FAST sensors placed on mice without tumors. (n=5 animal replicates in grey; Black Line = Average). (B) The relative resistance change of FAST sensors on mice with treated and untreated A20 tumors is statistically significantly different from the sensor drift experienced on mice without any tumors after 12 hours (One-way ANOVA with Tukey’s multiple comparison test; Line = Average).

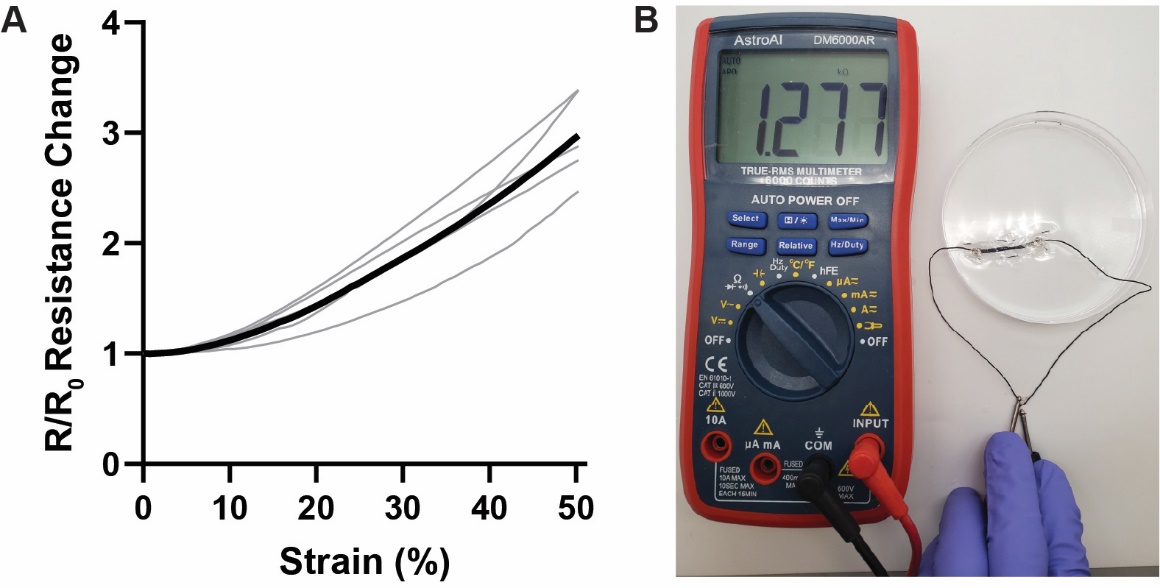

**Supplementary Figure S6: Encapsulated PDMS Strain Sensor Characterization.** (A) Strain versus resistance curves for a cracked gold PDMS strain sensor with an initial length of 23 mm. (B) Even when submerged in phosphate buffered saline, the resistance readouts from the encapsulated sensor are unaffected. (Individual curves. Bold line = Average).

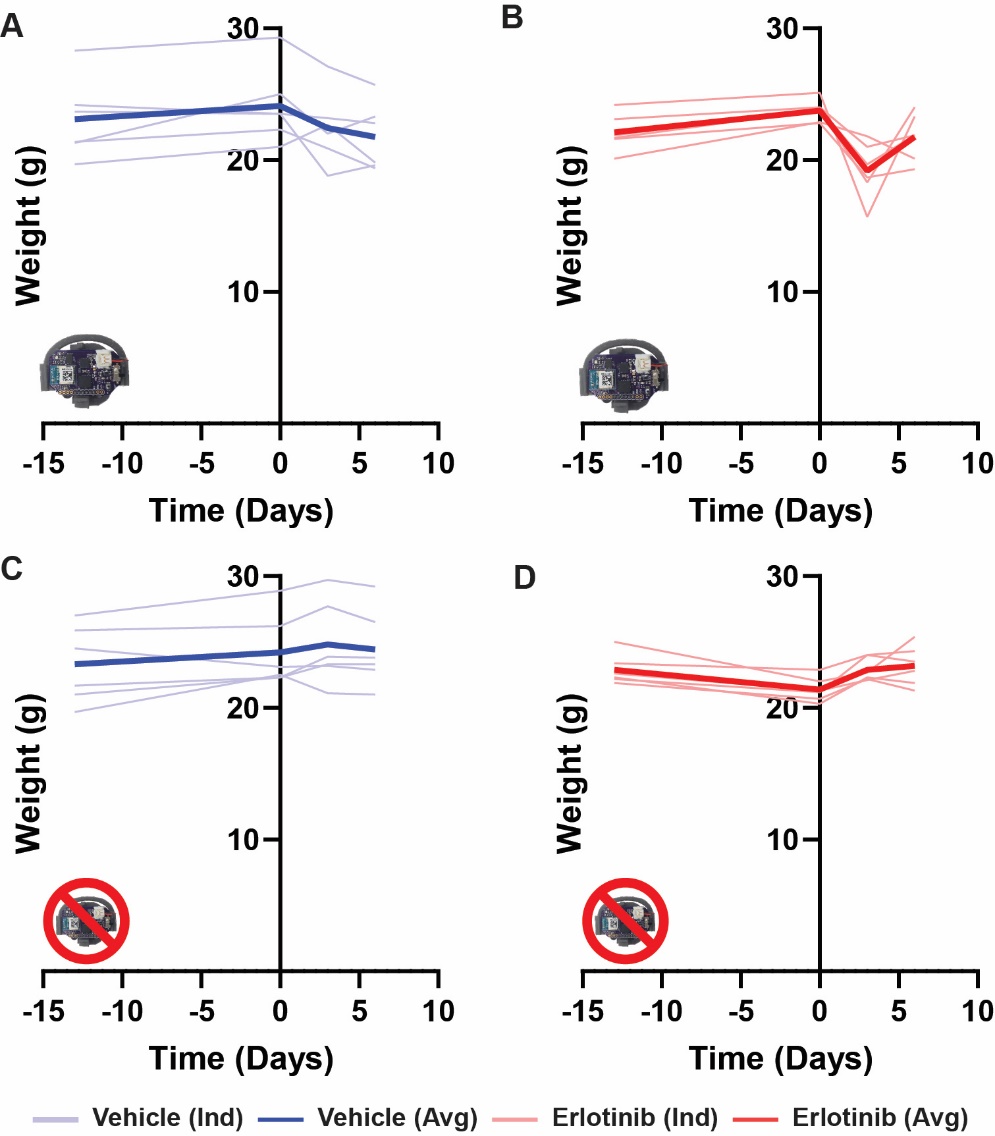

**Supplementary Figure S7: Mouse weight for HCC827 model over treatment period.** (A-B) Mouse weight over time for sensor-wearing (A) vehicle treated and (B) erlotinib treated mice. (C-D) Mouse weight over time for non-sensor-wearing (C) vehicle treated and (D) erlotinib treated mice. (Individual curves. Bold line = Average).

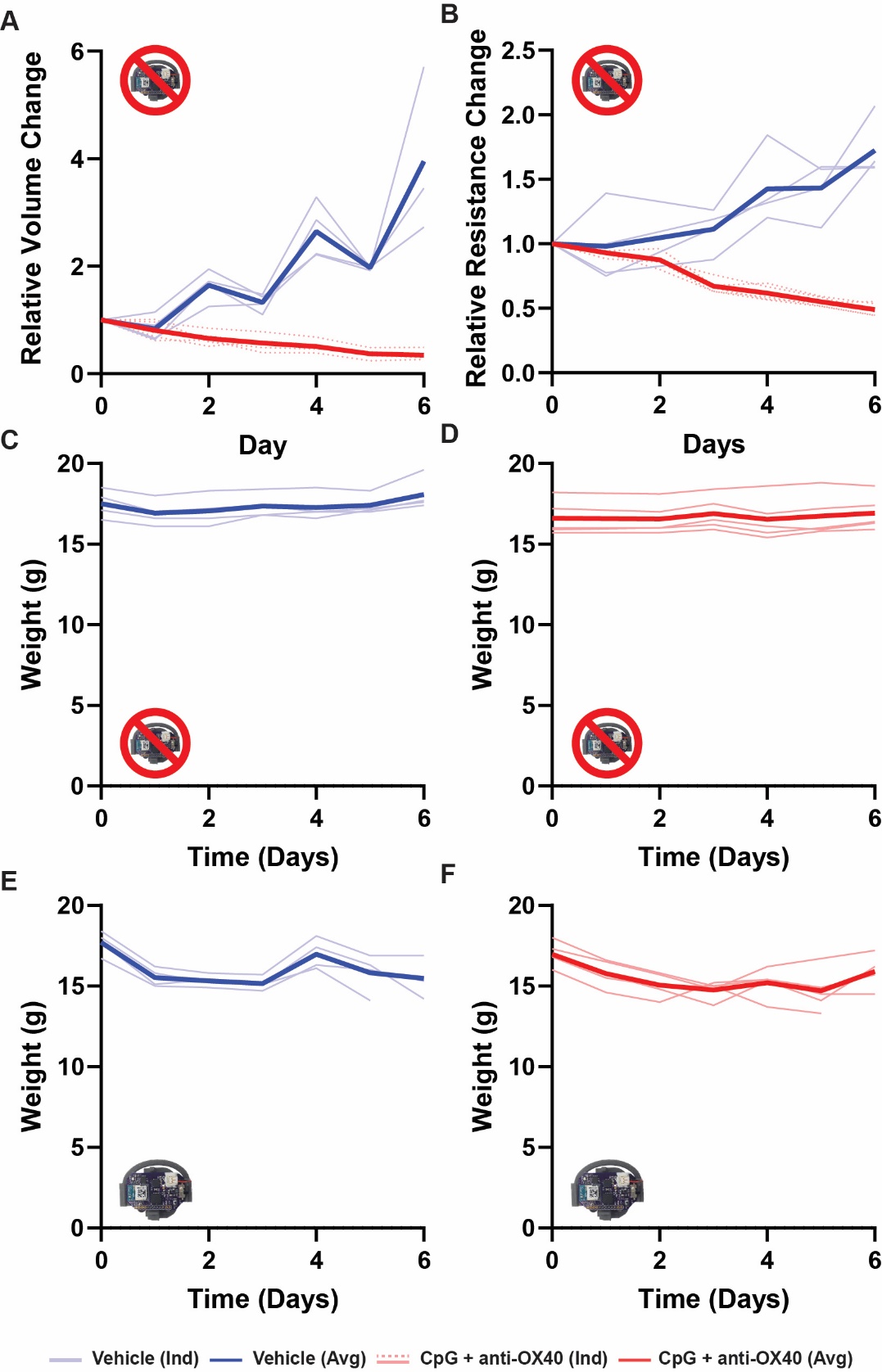

**Supplementary Figure S8: Mouse weight and controls for A20 model over treatment period.** (A-B) Relative tumor volume change and (B) Relative Resistance change from periodic sensor measurements for vehicle and CpG+aOX40 treated A20 tumors in mice that did not wear the continuous sensor. (C-F) Mouse weight over time for sensor-wearing (C) vehicle treated and (D) CpG + anti-OX40 treated mice without and (E,F) with continuous sensor wrap. (Individual curves. Bold line = Average. Vehicle n=4, Treatment n=5).

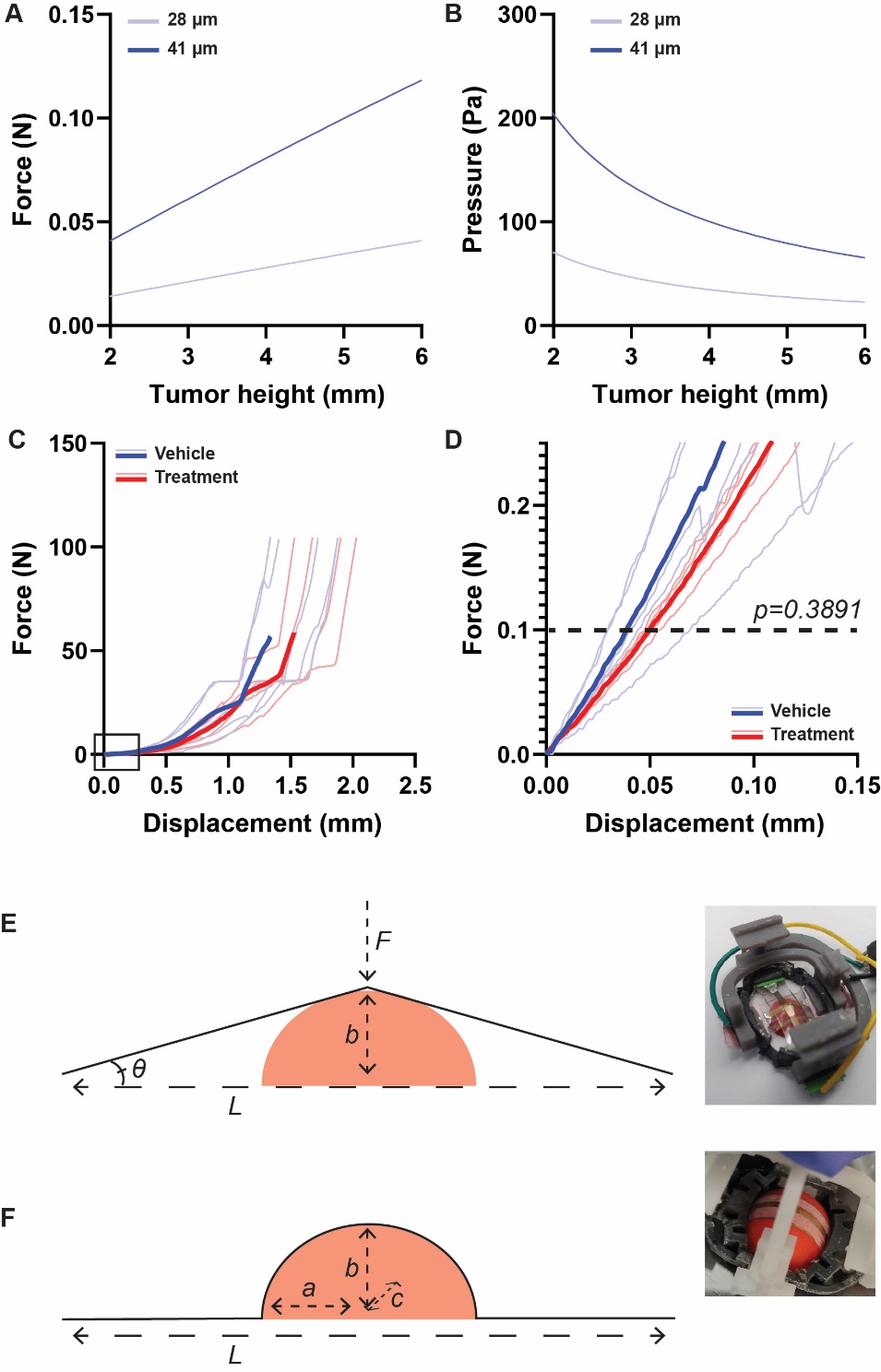

**Supplementary Figure S9: Estimates of force and pressure on tumor from sensor.** (A-B) Estimates for the (A) force and (B) pressure applied by the elastic SEBS sensor on the tumor. The pressure calculation assumes a tumor in which the length and width are 4x longer than the height. (C-D) Compression tests on Instron of vehicle treated and CpG+aOX40 treated A20 tumors. There is no statistically significant difference between the displacement values at a force of 0.1 N. (Unpaired Students t-test, n=5 Vehicle, n=4 treated, Individual lines are individual tumors from different mice, Bold Line=Average). (E-F) The sensor can wrap around the tumor in one of two ways, as denoted in the diagrams and images. The bottom design utilizes flexible bands built into FAST backpacks that enable the stretchable sensor to hug the tumor, increasing the total strain on the sensor. The diagram shown in panel C denotes the distances and angles used for the calculations of force and pressure.
